# Experimental warming reduces gut prokaryotic diversity, survival and thermal tolerance of the eastern subterranean termite, *Reticulitermes flavipes* (Kollar)

**DOI:** 10.1101/2020.09.19.300780

**Authors:** Rachel A. Arango, Sean D. Schoville, Cameron R. Currie, Camila Carlos-Shanley

## Abstract

Understanding the effects of environmental disturbances on the health and physiology of insects is crucial in predicting the impact of climate change on their distribution, abundance, and ecology. As microbial symbionts have been shown to play an integral role in a diversity of functions within the insect host, research examining how organisms adapt to environmental fluctuations should include their associated microbiota. Previous studies have shown that temperature affects the diversity of protists in termite gut, but less is known about the bacterial symbionts. In this study, subterranean termites (*Reticulitermes flavipes* (Kollar)) were exposed to three different temperature treatments characterized as low (15 °C), medium (27 °C), and high (35 °C). Results showed low temperature exposed termites had significantly lower CTmin and significantly higher SCP values compared to termites from medium or high temperature groups. This suggests that pre-exposure to cold allowed termites to stay active longer in decreasing temperatures but caused termites to freeze at higher temperatures. High temperature exposure had the most deleterious effects on termites with a significant reduction in termite survival as well as reduced ability to withstand cold stress. The microbial community of high temperature exposed termites showed a reduction in bacterial richness and decreased relative abundance of Spirochaetes, Elusimicrobia, and methanogenic Euryarchaeota. Our results indicate a potential link between gut bacterial symbionts and termite’s physiological response to environmental changes and highlight the need to consider microbial symbionts in studies relating to insect thermosensitivity.

## 1. Introduction

Temperature is considered a major factor influencing the distribution of many, if not all, living organisms (Doucet et al. 2009). As poikilotherms, insects are particularly susceptible to fluctuations in temperature as their internal body temperature is directly related to external environmental conditions. Current predictions of global climate change suggest insects will be exposed to more extreme temperatures with increased frequency, which will directly impact seasonality, distribution, survival, and overall fitness of both beneficial and pestiferous insect species in the future (Wernegreen 2012, Emerson 1955, Lee and Denlinger 1985, Smith and Rust 1994, Worland et al. 1998, Sinclair 2001, Slabber et al. 2007, Bonnett et al. 2012, Sinclair et al. 2015, Alford et al. 2018). Thus, studies examining temperature tolerance and the various mechanisms that contribute to this tolerance are becoming increasingly important.

The level of vulnerability to fluctuations in environmental conditions (e.g. increased exposure to thermal stress) for various insect species is thought to be linked with the underlying phenotypic plasticity present in these organisms. Researchers have highlighted the diversity of mechanisms used by insects to acclimate to novel environments or shifts in environmental conditions. Examples include behavioral modifications (e.g. habitat selection, temporal activity), physiological changes (e.g. production of enzymes, antioxidants, proteins, lipids, carbohydrates, trehalose, etc.), and/or phenotypic alterations (e.g. reproduction, diapause, seasonal phenology) (Kingsolver and Huey 1998, Huey et al. 2003, Danks 2005, Clark and Worland 2008, Huey 2010, Lacey et al. 2010). A number of studies have also identified insect-associated microorganisms that help facilitate thermal tolerance such as the presence of ice nucleating bacteria which serve to regulate freezing in preparation for overwintering in certain freeze-tolerant insects (Lee et al. 1991, Lee et al. 1993, Wernegreen 2012, Moghadam et al. 2018). Ferguson et al (2018) found the composition of the gut microbiome to shift seasonally in crickets (*Gryllus veletis*) which increased freeze tolerance and immunity.

As studies related to microorganisms and thermal tolerance continue to expand, research has shown the interactions between the microbial community and host fitness to be complex. For example, while associations with certain bacteria have the potential to enhance thermal tolerance, it has also been suggested that primary symbionts of certain insect species (i.e. microorganisms that perform specific, essential functions that influence host fitness), can actually constrain, rather than facilitate, thermal tolerance (Wernegreen 2012). Various researchers have hypothesized that because symbionts have co-evolved with their host, they are physiologically and genetically constrained, and thus more susceptible to environmental fluctuations (Wernegreen 2012). Bestion et al. (2017) reported a 34% loss in microbial diversity and shifts in the relative abundance of major bacterial phyla in the gut of the lizard, *Zootoca vivipara*, after only a 2-3 °C increase in temperature. Prado et al. (2010) showed a significant reduction in bacterial symbionts in the guts of two stinkbug species after high temperature exposure (30 °C). Additionally, Montllor et al. (2002) found guts of heat-stressed pea aphids had a reduced number of bacteriocytes which house their obligate primary symbiont, *Buchnera*.

Perhaps one of the most studied insect-symbiont relationships is that of wood-feeding termites and their gut symbionts, which are essential for breakdown of lignocellulosic materials (Breznak and Brune 1994, Radek 1999, Brune 2014). In North America, subterranean termites in the family Rhinotermitidae are responsible for the majority of economic costs associated with damage to wooden products and structures (Su and Scheffrahn 2000). Members of Rhinotermitidae are broadly distributed in both temperate and tropical regions, although individual termite species and populations tend to thrive within relatively narrow temperature ranges and under specific environmental conditions (Esenther 1969, Davis and Kamble 1994). As microbial symbionts have been credited as significant drivers of species evolution and diversification in termites (Mueller et al. 2011), there is the potential for termite-associated microorganisms to influence thermal tolerance plasticity and constrain termite geographical distributions.

A number of researchers dating back to the 1920s have examined the effect of temperature on termites (Esenther 1969, Haverty and Nutting 1974, Sponsler and Appel 1991, Davis and Kamble 1994, Cabrera and Kamble 2001, Hu and Appel 2004, Hu and Song 2007, Gautam and Henderson 2011), as well as on their symbiotic gut protozoa (Cleveland 1923 & 1924, Cook and Smith 1942, Mannesmann 1969, Smythe and Williams 1972, Belitz and Waller 1998, Cabrera and Kamble 2004). Cleveland (1924) showed that all gut protozoa were killed after 24-hour incubation at 36 °C in *Reticulitermes flavipes* (Kollar). Smythe and Williams (1972) observed no effect on protozoa at 29 °C and report the highest tolerable temperature range for *R. flavipes* survival to be between 31.5 °C and 33 °C. Despite this abundance of research relating to the effect of temperature on protozoa, the effect on gut bacteria has been mostly ignored. One exception to this is a study of termite cold tolerance, where Cabrera and Kamble (2004) found the supercooling point to be the lowest in *R. flavipes* workers treated with antibiotics, and suggested that symbionts (either protists or bacteria) may be acting as ice nucleators in the termite hindgut.

The primary objectives of this study were to: 1) examine the physiological performance of eastern subterranean termites, *R. flavipes*, after prolonged exposure to three different temperature conditions using measurements of feeding activity, mortality, and cold tolerance (supercooling point (SCP) and critical thermal minimum (CTmin)), 2) evaluate how an increase or decrease in temperature affects the microbial community of the termite gut and associated nest (i.e. soil material used to construct galleries), and 3) examine how these microbial shifts relate to thermal tolerance. These objectives aim to gain a better understanding of temperature induced physiological and microbiological shifts of the termite gut microbial community and their potential functional role in termite thermal tolerance.

## 2. Materials and Methods

### 2.1 Experiment Set-Up and Temperature Treatment

Termite experiments were performed in sterile plastic petri dishes containing sterile soil (10 g), and sterile DI water (1 mL). A southern yellow pine wood block (*Pinus spp*.) (40 × 25 × 2 mm) which was conditioned at 27 °C/30% relative humidity (RH), weighed and autoclave-sterilized, was included in each dish as a food source. A total of 100 *R. flavipes* workers (∼3^rd^ – 4^th^instar), originally collected from Janesville, Wisconsin and maintained under laboratory conditions (27 °C, 80% RH), were added to each dish. Dishes were then sealed with parafilm to maintain moisture levels. Termites were exposed for 4-weeks to one of three temperatures, characterized as low (15 °C), medium (27 °C), or high (35 °C), with 5 replicate dishes per temperature treatment. These temperatures were selected to represent a range of environmental challenges and were within the experimental ranges used by previous researchers that elicited physiological effects in termites.

### 2.2 Termite Guts and Soil Samples

After 4-weeks, soil material (0.25 g) manipulated by the termites (i.e. soil pasted to the side of the container by termites using salivary secretions and faeces) was sterilely collected in duplicate from each of the petri dishes into sterile 2 mL microcentrifuge tubes. Pine wood feeder blocks from each dish were brushed free of debris, re-conditioned to a constant weight and re-weighed to determine the amount of termite feeding at each temperature treatment. Termites from each dish were counted to determine mortality. Termites were then randomly selected from each temperature group for use in cold tolerance assays (31 termites/test group), or for dilution series to determine bacterial abundance (5 termites/replicates/treatment). Another subset of termites (n=20 per treatment group) were sacrificed by freezing, rinsed in 70% EtOH to sterilize the termites externally, and their guts were extracted for 16S rRNA amplicon sequencing. Extracted guts from termites collected directly from a field site in Janesville, Wisconsin at 11 time points across late spring, summer and early winter seasons were also included for comparative purposes.

### 2.3 DNA Extraction and 16S rRNA Amplicon Sequencing

Termite guts were extracted directly into ZR bashing bead lysis tubes (Zymo Research, Irvine, CA) and a mix of 600 µl of tissue and cell lysis solution and 2 µl Proteinase K from the MasterPure Complete DNA and RNA Purification Kit (Epicentre-Lucigen, Middleton, WI) was added to each sample tube. All tubes were then vortexed vigorously for 2 minutes. Hereafter, DNA extraction methods followed the Epicentre-Lucigen total nucleic acids purification protocol. DNA was extracted from 0.25 g of collected soil material using the PowerSoil DNA Isolation Kit (MoBio, Carlsbad, CA) following the provided protocol. Both termite gut and soil DNA was quantified using the Qubit Fluorometer (HS-assay kit; Invitrogen, Carlsbad, CA) and diluted to 20 ng/µl.

Polymerase chain reaction (PCR) was done using tagged MiSeq primers targeting the V4 region of the 16S rRNA gene [primers: forward -GTGCCAGCMGCCGCGGTAA; reverse -GGACTACHVGGGTWTCTAAT] (Kozich et al. 2013). Reaction mixtures for each sample (24 µl) included 12 µl KAPA HiFi HotStart ReadyMix (2X) (KAPA Biosystems, Boston, Massachusetts), 1.5 µl NanoPure water, 1 µl forward primer (10 µM), 1 µl reverse primer (10 µM), and 8 µl diluted DNA (20 ng/µl). All samples were run in triplicate under the following conditions: initial denaturation at 95 °C for 3 minutes, 25 cycles of 95 °C for 30 seconds, 55 °C for 30 seconds, 72 °C for 30 seconds, and a final elongation step at 72 °C for 5 minutes. A non-template control reaction was included and submitted for sequencing.

Amplicon libraries were sequenced with the paired-end Illumina MiSeq platform at the University of Wisconsin-Madison Biotechnology Center. Reads were preprocessed, assembled, aligned, and classified using the mothur pipeline (Schloss et al. 2009). Classification was based on the Silva SEED release 132 database. Operational taxonomic units (OTUs) were defined using a 99% similarity threshold. Z-score normalized data was calculated as Z= (x-u)/d, where x is the relative abundance of a taxa, u is the mean relative abundance of a taxa across all the samples and, d is the standard deviation across all the samples. A phylogenetic tree of representative sequences of the top OTUs across all samples was performed in the MEGA v7 using the maximum likelihood method with 500 bootstrap replicates, assuming a General Time Reversible (GTR) substitution model. Richness of OTUs was estimated using Chao-1 index (Chao and Chiu 2016), and non-metric multidimensional scaling (nMDS) and permutational analysis of variance (PERMANOVA) were used to quantify differences among samples and assess statistical significance, using PRIMER-e v 7.0.13.

### 2.4 Thermal Tolerance Tests

Termites used for this study originated from a population of *R. flavipes* along the northern range of its distribution where they are likely limited by cold seasonal conditions. Thus, two cold tolerance assays, supercooling point and critical thermal minimum, were selected as the physiological measures of thermal tolerance.

#### 2.4.1 Supercooling Point (SCP)

Two worker termites (3^rd^ or 4^th^ instar) were placed into each of eight 1.5 ml microcentrifuge tubes (n=16 per treatment group). T-type thermocouples were wrapped near the base with cotton rounds and pushed in, next to the termites at the bottom of the tube to measure termite temperature. The other end of the thermocouple was inserted into an 8-channel TC-08 Data Logger (Pico Technology, Tyler, TX) set to record temperature of all 8 channels, at a sampling interval of 2 seconds. Tubes were then inserted into a floating tube rack and carefully placed into the refrigerated water bath (Grant Instruments, Cambridge, United Kingdom) containing 50:50 propylene glycol to water. The water bath was controlled using an external circulating pump and accompanying LabWise software, which was programed for an initial 10-minute acclimation step at 20 °C followed by a decrease in temperature at a rate of 0.5 °C/minutes down to 5 °C, then at 0.2 °C/minute until it reached −15 °C. Measurements of SCP were determined by a spike in temperature as measured by the data logger, representing the temperature at which freezing is initiated.

#### 2.4.2 Critical Thermal Minimum (CTmin)

A custom aluminum cooling block was connected to a refrigerated water bath (Grant Instruments, Cambridge, United Kingdom) with insulated plastic tubes, allowing for cycling of the 50:50 propylene glycol to water between the water bath and aluminum block. Water bath temperature was controlled using an external circulating pump and accompanying LabWise software, programed for an initial 10-minute acclimation step at 20 °C followed by decreasing temperatures at a rate of 0.2 °C/minute down to −15 °C. Groups of 3 termite workers (3^rd^ or 4 ^th^instar) were added to 5 of the 6 small culture plates attached to the center of the aluminum cooling block with thermal conducting tape (n=15 per treatment group). Each culture plate had a small hole drilled in the lid to prevent condensation and so that a T-type thermocouple could be inserted for accurate temperature measurements. Temperature was recorded using a TC-08 Data Logger (Pico Technology, Tyler, TX) in one of the six culture plates. Measurements of CTmin were determined as the temperature at which termites displayed a lack of righting ability after being gently flipped over with a paint brush, indicating that they could no longer coordinate their muscle movements. Once CTmin was reached, the termite was removed from the test dish into a 12-well plate and allowed to recover at room temperature (Note: if termites did not recover, the CTmin threshold was passed and the data from that termite was deemed invalid).

## 3. Results and Discussion

### 3.1 Effect on the Termite

#### 3.1.1 Survival and Feeding

During the 4-week test, termites in the low temperature group consumed significantly less of the pine wood feeder block compared to termites in the other two temperature treatment groups (Fig. 1a). Termite survival was comparable between the low and medium temperature groups but was significantly lower in termites exposed to the high temperature treatment (Fig. 1b). These results suggest that while termites ate less at lower temperatures, this did not negatively impact survival. Termites in the high temperature group consumed slightly less wood material than the medium temperature termites, which was likely the result of increased mortality. Since wood consumption was not significantly lower however, it is likely that termite mortality occurred towards the end of the test period.

**Figure 1.**
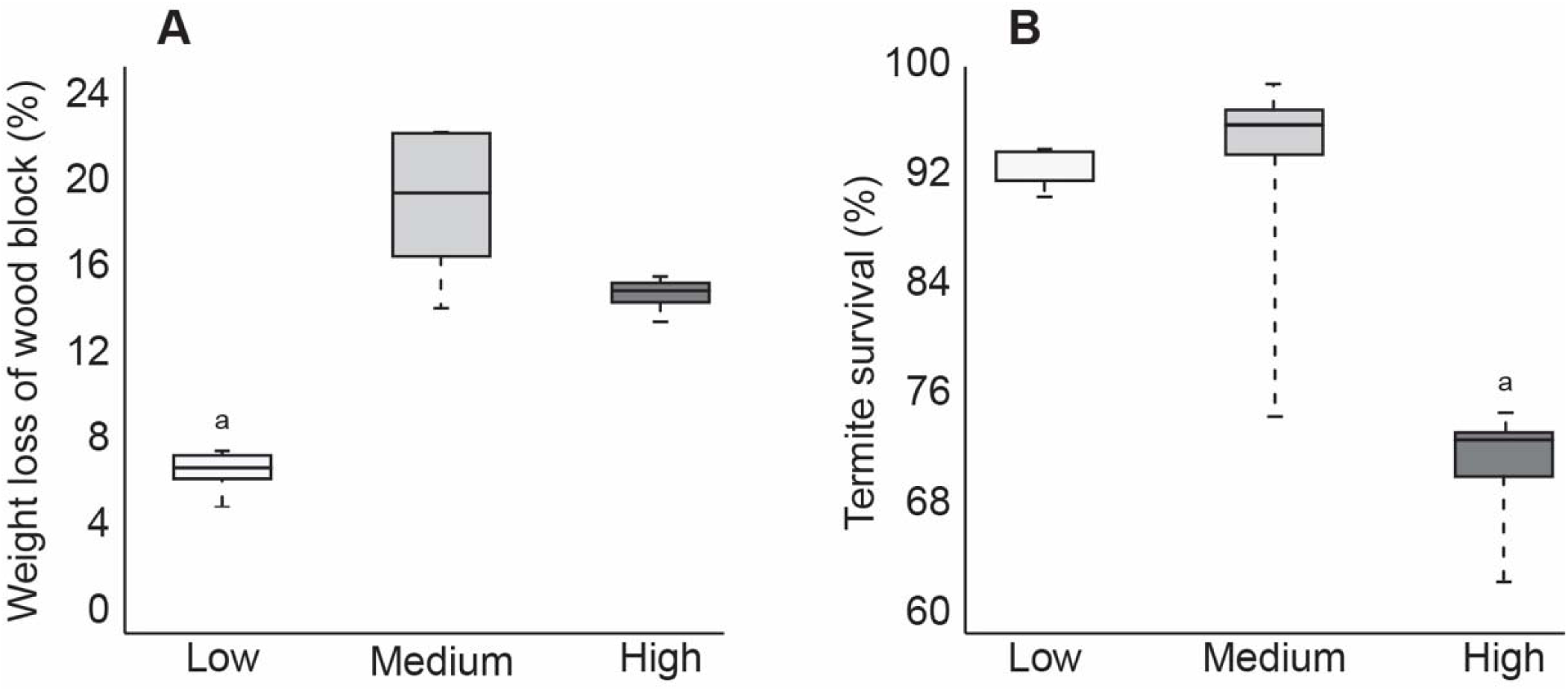
Boxplot showing average amount of feeding (a) and average termite survival (b) after four weeks at low, medium, or high temperatures (p < 0.05).

#### 3.1.2 Cold Tolerance

Data from cold tolerance tests, supercooling point (SCP) and critical thermal minimum (CTmin) are shown in Figure 2. These tests showed a significantly higher CTmin (mean 15.6 °C) in high temperature exposed termites, suggesting increased susceptibility to decreases in temperature. Termites acclimated at low temperature had a significantly lower CTmin (mean 5.69 °C), but a significant increase in SCP (mean −5.9 °C) compared to termites acclimated at the medium (mean CTmin 7.3 °C/ SCP −8.4 °C) or high (mean SCP −7.6 °C) temperatures. This suggests that pre-exposure to cold allowed termites to stay active longer in decreasing temperatures but caused termites to freeze more easily. Although the higher SCP suggests an increase in freezing susceptibility, this is not altogether unexpected as termites are thought to be freeze-intolerant/freeze avoidant insects (Mail 1930, Esenther 1969, Cabrera and Kamble 2001 & 2004, Hu and Song 2007, Clarke et al. 2013) and SCP represents a lower limit of survival that may be dependent on multiple physiological factors (Renault et al. 2002, Lee 2010). Results from this study are supported by those from a similar study of *R. flavipes* cold tolerance where they showed an increase in SCP after pre-exposure to 10 °C, but concluded that lowering supercooling point is not likely a factor in cold acclimation as termites are not freeze tolerant insects (Davis and Kamble 1994). While our results reinforce this conclusion, the reason for higher SCP values after cold acclimation remains unknown but may relate to the abundance of ice-nucleating agents in the termite gut. Other freeze-intolerant insect species have been shown to reduce ice nucleating bacteria and/or microbially produced compounds (e.g. calcium carbonate, potassium phosphate, uric acid, certain amino acids, proteins, steroids) which lead to freezing injury (Vonnegut 1949, Head 1961, Strong-Gunderson et al. 1990, Kawahara 2002, Clark and Worland 2008, Lee 2010). When ice nucleators are present, they promote freezing at higher temperatures, which serves to limit ice formation to extracellular fluids in freeze tolerant species (Neven et al. 1989, Duman et al. 2010). Thus, high SCP in low temperature groups might be indicative of the presence of ice nucleating microorganisms that either increased during the cold acclimation period or are normally excreted seasonally in field termites in preparation for winter. Accordingly, lower SCP values in the other two temperature groups may be the result of reduction or shift in gut microbiota that would otherwise act as ice nucleating agents. Cabrera and Kamble (2004) showed antibiotic-treated *R. flavipes* workers to have the lowest SCP values and suggest this may be the result of removing ice-nucleating microorganisms. The significance of these results remains to be determined but could hint at one possible transition by which freeze intolerant insects, such as termites, may evolve to become freeze tolerant.

**Figure 2.**
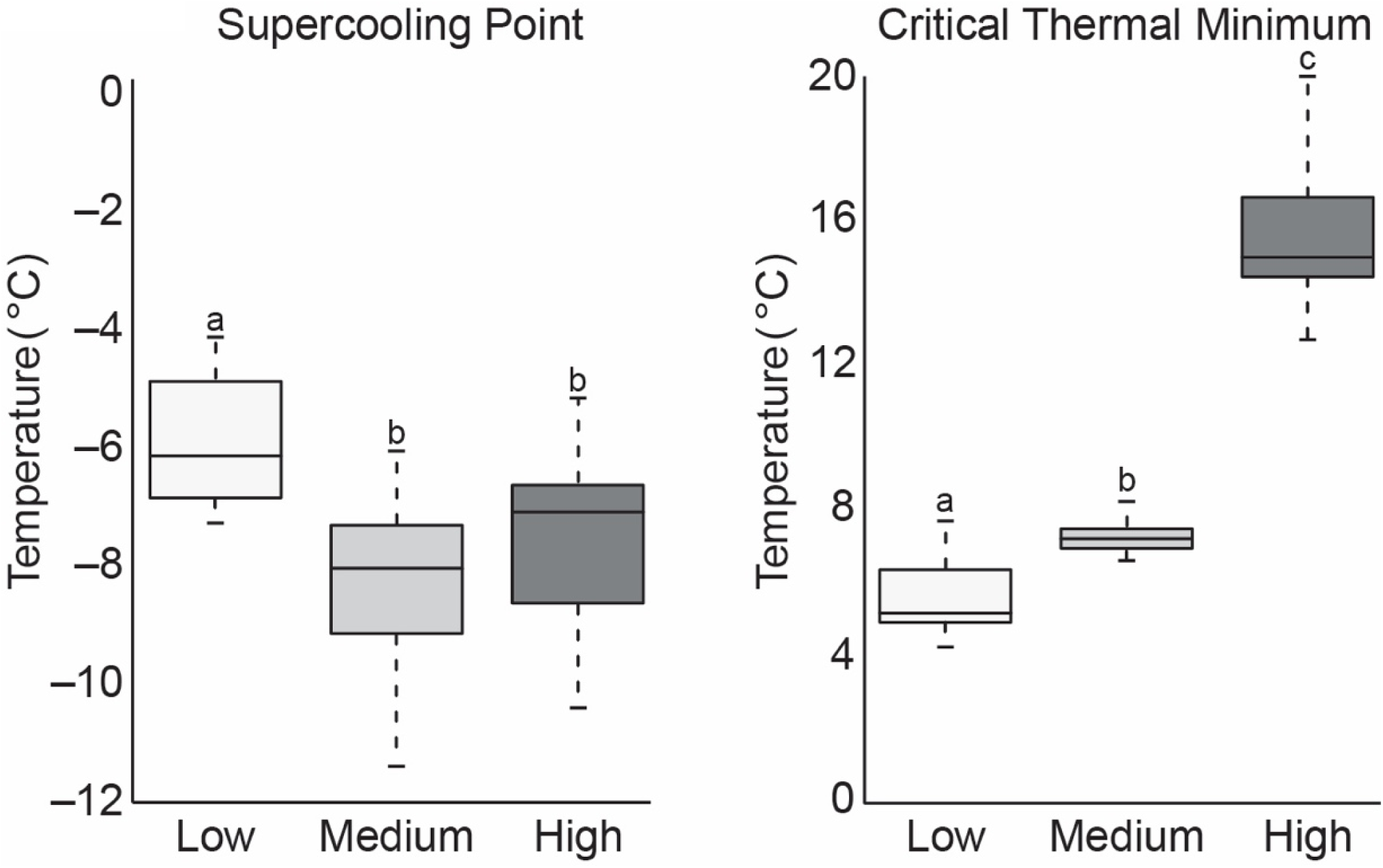
Boxplot of SCP and CTmin values of termite workers after four weeks of exposure to low, medium, or high temperatures (treatments that do not share the same letter are significantly different p ≤ 0.05).

### 3.2 Effect on the Termite Gut

#### 3.2.1 Similarity and Diversity: Gut Microbial Community

The relative abundance of bacterial phyla from each of the test groups and from field collected gut samples are shown in Figure 3. The microbial community of field collected *R. flavipes* guts were dominated by Spirochaetes, followed by Firmicutes, Bacteroidetes, Elusimicrobia, and Proteobacteria, which is comparable to results from other studies that characterize the core microbiota of *Reticulitermes* spp. guts (Ohkuma and Brune 2011, Boucias et al. 2013, Brune 2014, Benjamino and Graf 2016). Within the temperature treatment groups, the microbial community of guts from the medium temperature groups most closely resembled those of field populations at the phyla level although they did show a small reduction in the relative abundance of Spirochaetes and an increase in Planctomycetes.

**Figure 3.**
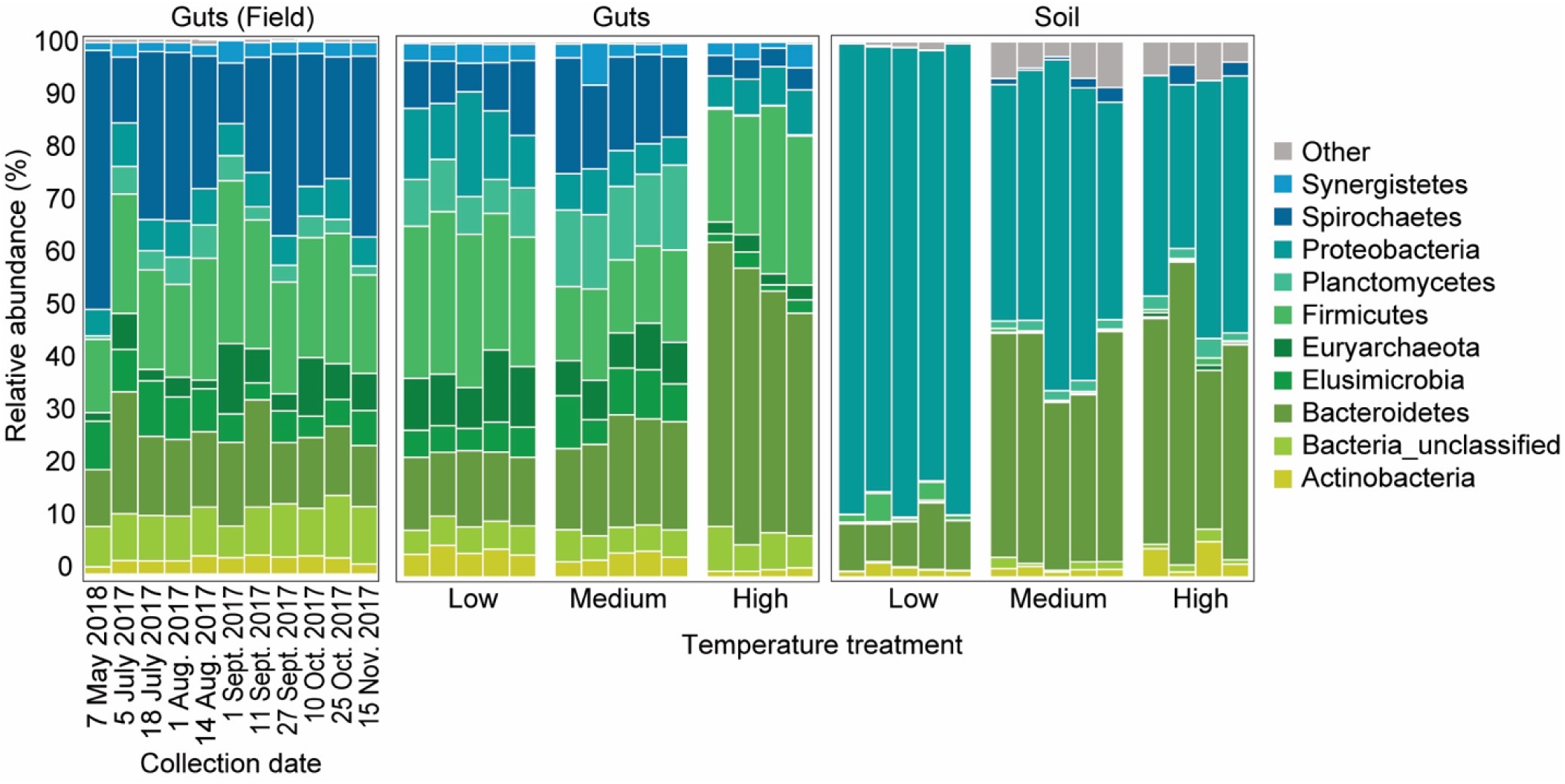
Average relative abundance (%) of bacterial phyla from field collected termite guts (left) and termite guts and soil material after low, medium, or high temperature exposure (right).

The gut microbial community composition was significantly more similar within treatments than within treatments (Table 1 and 2, Fig. 4b). However, termite guts from the medium and low temperature groups showed more similarity to each other, and to those collected directly from the field, than termite guts from the high temperature group. Additionally, Chao-1 richness was significantly reduced in guts from the high temperature group compared to those from field samples or from the other two temperature groups (Fig. 5). Other studies have also shown the gut of poikilotherms to have lower bacterial diversity after exposure to heat stress (Bestion et al. 2017, Fontaine et al. 2018). Studies in bivalves (Abele et al. 2002) and lepidopterans (Cui et al. 2011) have shown exposure to thermal stress can lead to an increase in the production of reactive oxygen species. One possible explanation for the observed lower bacterial diversity under high temperatures is that different species of gut symbionts show different levels of tolerance to reactive oxygen species produced by the host, which could result in changes in microbiome structure (Obata et al. 2018). More studies are needed to test this hypothesis. Moreover, our results show that temperature shapes not only termite–protist symbiosis (Cleveland, 1924; Belitz &Waller, 1998), but also the bacterial members of this complex symbiosis.

**Table 1.** Average Bray-Curtis similarity (%) of the bacterial communities within and between groups.

| Average similarity | Treatment | Soil | Guts |
| --- | --- | --- | --- |
| Within | Field | - | 58.4 |
|  | Low | 42.9 | 63.7 |
|  | Medium | 52.2 | 64.7 |
|  | High | 49.0 | 64.6 |
| Between | Field-Low | - | 50.2 |
|  | Field-Medium | - | 50.0 |
|  | Field-High | - | 30.4 |
|  | Low-Medium | 27.5 | 57.2 |
|  | Low-High | 25.0 | 34.7 |
|  | Medium-High | 46.7 | 40.0 |

**Table 2.** Pair-wise PERMANOVA results for Bray-Curtis similarities of the bacterial communities from the termite guts and termite-manipulated soil materials after temperature treatment (low, medium, or high) and from field collected samples.

|  | Comparison groups | t | P-value |
| --- | --- | --- | --- |
| Termite guts | Field, High | 3.5506 | 0.001 |
|  | Field, Low | 2.1921 | 0.002 |
|  | Field, Medium | 2.2398 | 0.002 |
|  | High, Low | 3.3492 | 0.01 |
|  | High, Medium | 3.0541 | 0.006 |
|  | Low, Medium | 1.7686 | 0.008 |
| Soil | High, Low | 2.2329 | 0.009 |
|  | High, Medium | 1.3191 | 0.006 |
|  | Low, Medium | 2.3433 | 0.008 |

**Figure 4.**
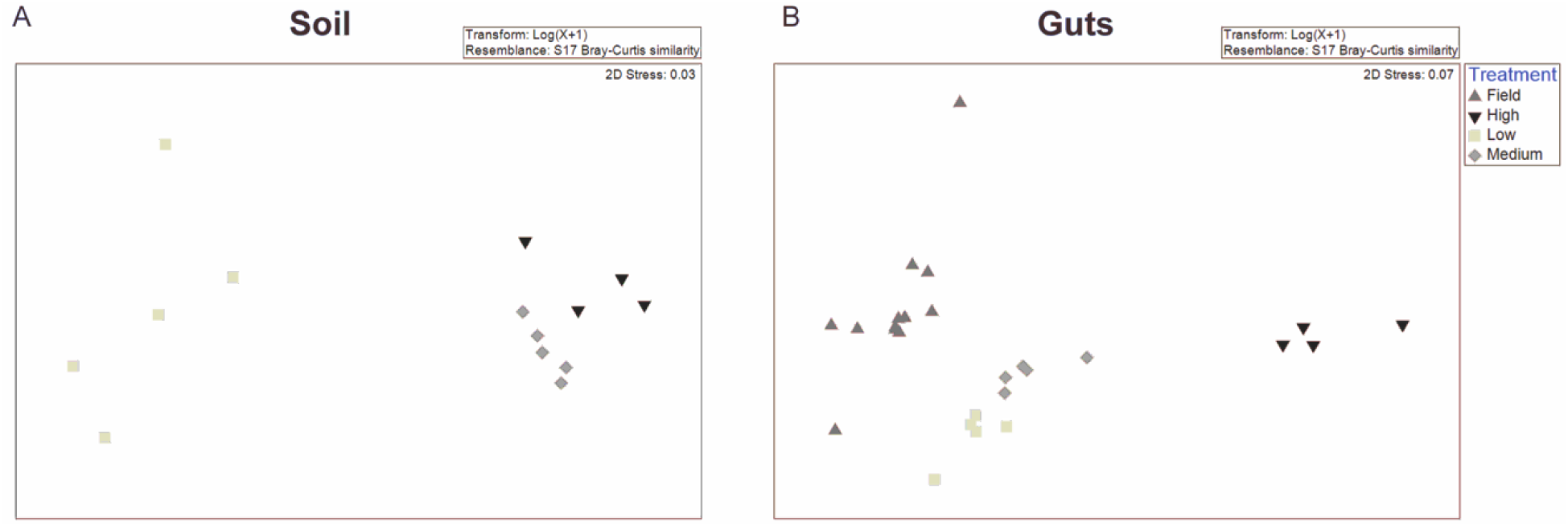
Non-metric Multidimensional Scaling (NMDS) plot based on the Bray-Curtis distance of the bacterial communities of (a)termite-manipulated soil materials after temperature treatment (low, medium, or high) and (b) of termite guts after temperature treatment (low, medium, or high) and from field collected samples.

**Figure 5.**
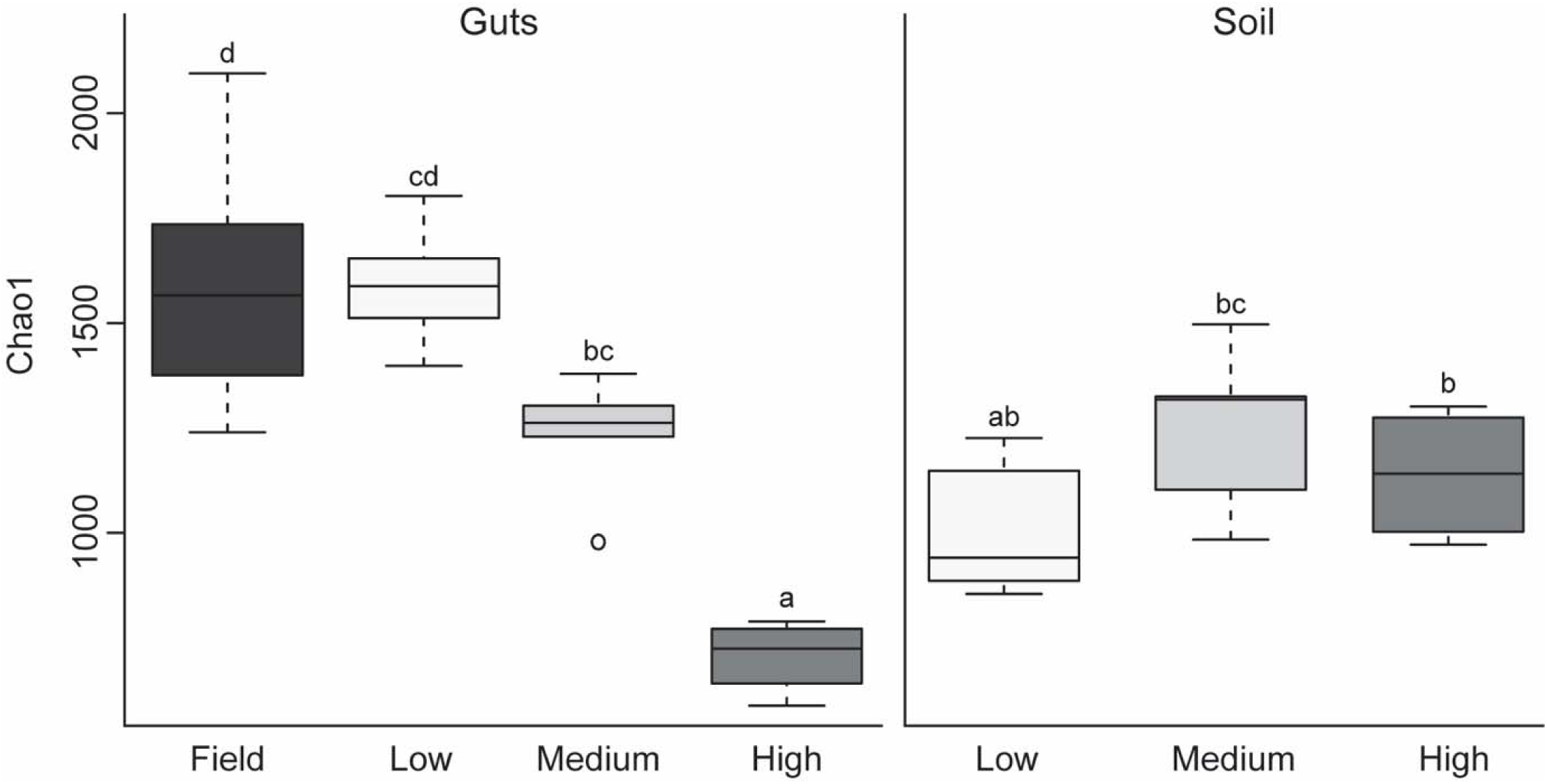
Boxplot showing estimated microbial diversity based on Chao-1 for field collected samples and termites exposed to low, medium, or high temperatures (treatments that do not share the same letter are significantly different p ≤ 0.05).

In this study, high temperature treatment caused the most dramatic shift in the overall microbial community of the termite gut, specifically an increased relative abundance of Bacteroidetes to nearly 50% of the gut community composition (Table 1, Fig. 3). Examination of the top 20 OTUs from guts samples of all temperature treatment groups, showed four OTUs to be members of Bacteroidetes, two of which belong to Dysgonomonadaceae (OTUs: 00020, 00005). Members of this family have recently been shown to be ectosymbionts on intestinal nematodes in wood-feeding cockroaches (Murakami et al. 2019), although their significance in this study remains to be determined.

High temperature treated groups also showed reduced relative abundance of Spirochaetes, Elusimicrobia, and Euryarchaeota. These shifts in prokaryote communities may be connected, at least in part, to mortality of various protist species. Phylogenetically lower termites maintain a diversity of gut protists that are essential for breakdown of lignocellulosic materials in wood-feeding termites (Breznak and Brune 1994, Hongoh 2011, Brune 2014). The external surface of these flagellate protists has been shown to serve as an attachment surface for numerous gut bacteria while others have been shown to house bacterial endosymbionts (Brune 2014). Spirochaetes, for example, have been found to be associated with gut flagellates (Noda et al. 2003), although many can be found free within the hindgut fluid (Brune 2014). Within the Spirochaetes, species of *Treponema* represent members of the core microbial community in guts of *Reticulitermes* spp. where they are thought to either serve as CO_2_ reducing acetogens or contribute to fixation of atmospheric nitrogen (N_2_) (Noda et al. 2003, Benjamino and Graf 2016). Of the 20 most abundant OTUs from all groups, four were identified as *Treponema* spp. (OTUs: 00004, 00016, 00023, 00025) all of which were reduced in the high temperature treatment group (Fig. 6). Decreased relative abundance of specific OTUs after high temperature exposure was also observed in Elusimicrobia candidate genus *Endomicrobium* (OTU00022) which are known cytoplasmic protist endosymbionts (Stingl et al. 2005) and in a phylum of methanogenic archaea, Euryarchaeota *Methanobrevibacter* (OTU00008). Euryarchaeota is a phylum of methanogenic archaea that has been identified from the guts of all extant families of termites (Purdy 2007). These methanogens have mostly been found as ecto- and endosymbionts of termite gut protozoa although some have been shown to be associated with the gut wall (Ohkuma et al. 2006). Species of *Methanobrevibacter* specifically have been repeatedly identified from termite guts where they are suggested to utilize H_2_ and CO_2_ (Tokura et al. 2000). Thus, these data suggest that high temperatures likely have a negative impact termite gut protozoa, which agrees with results from Smythe and Williams (1972) who showed reductions in symbiotic protists (particularly those in the genera *Pyrsonympha* and *Dinenympha*) in guts of subterranean termites held at temperatures between 30 °C to 31.5 °C. These results support the hypothesis that the deleterious effects from high temperature exposure on termite survival may be indirectly related to the thermosensitivity of the microbial symbionts of the termite host.

**Figure 6.**
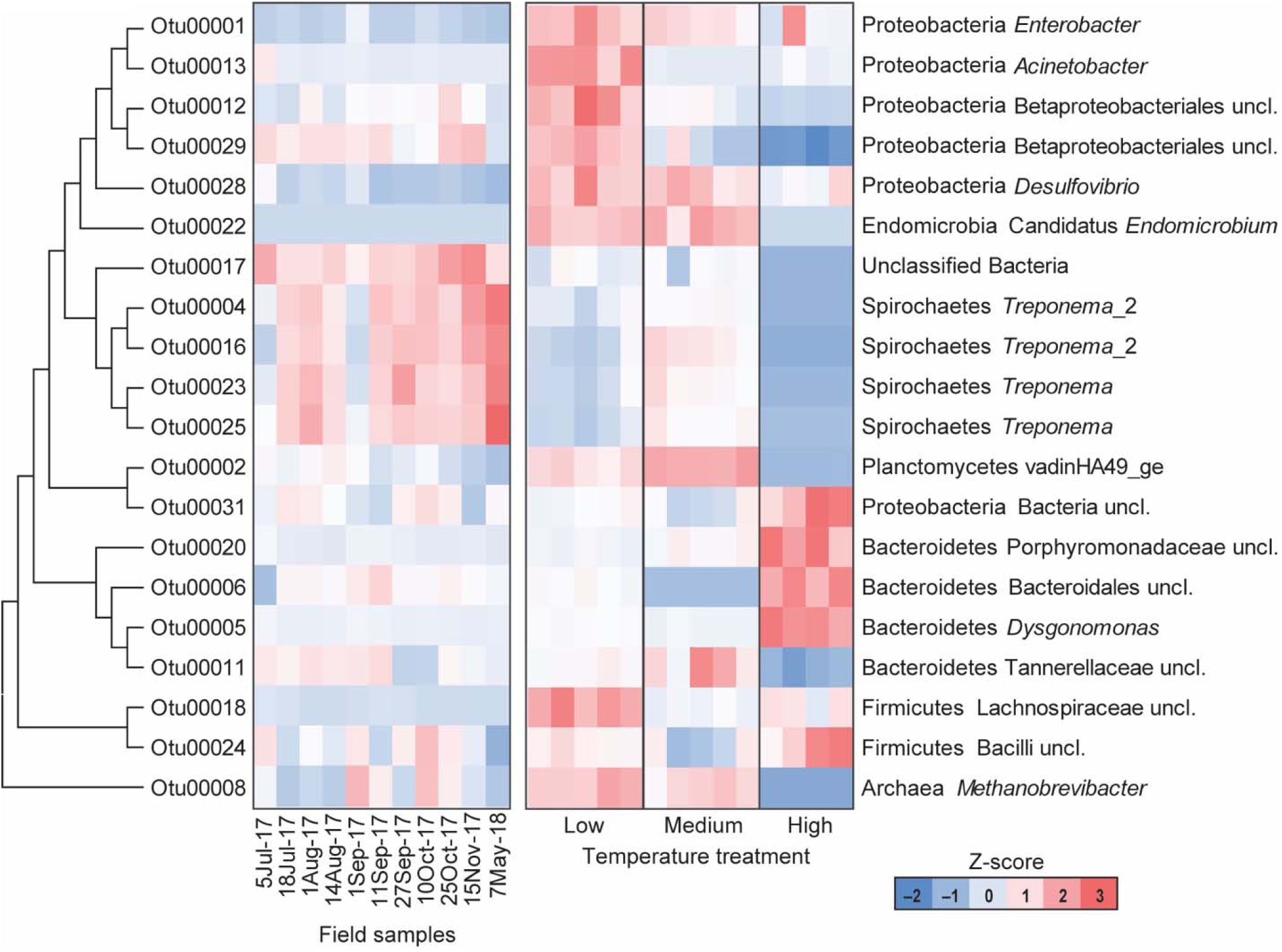
Heatmap representing the relative abundance of the top 20 OTUs from field collected termite guts (left) and termite guts after low, medium, or high temperature exposure (right).

Guts from low temperature treatment groups showed a reduction in the relative abundance of Spirochaetes and an increase in the relative abundance of Firmicutes, Lachnospiraceae (OTU 00018), in particular. Additionally, low temperature gut samples had nearly twice the relative abundance of Proteobacteria compared to the other temperature groups or field samples, specifically Acinetobacter (OTU00013), Desulfovibrio (OTU00028), Betaproteobacteriales (OTU00012, OTU00029), and *Enterobacter* (OUT00001) (Fig. 6). These data agree with a study examining seasonal shifts in the gut microbiome of spring field crickets (*Gryllus veletis*) that also showed an increase in the relative abundance of Proteobacteria after cold temperature exposure (Ferguson et al. 2018). This association of Proteobacteria and cold temperature exposure may correspond to the known ice nucleating activity of certain members of this phylum (e.g. *Pseudomonas* spp., *Enterobacter* spp., *Xanthomonas* spp.) (Lee et al. 1991, 1993, 1995). Therefore, the increased relative abundance of Proteobacteria in termite guts from the low temperature group may relate to cold tolerance, specifically the increase in SCP in termites pre-exposed to cold.

### 3.3 Effect on Termite-Manipulated Soil

#### 3.3.1 Similarity and Diversity: Soil Microbial Community

Samples from the low temperature group had the highest bacterial abundance, but also showed lower microbial diversity compared to the other temperature treatments based on Chao-1 (Fig. 5, Table 1). Comparisons of soil samples showed a distinct shift in the microbial community after low temperature treatment compared to the distinct, but more similar communities from the high and medium temperature samples (Fig. 4a, Table 1 and 2).

The relative abundance of bacterial phyla from termite-manipulated soil material is shown in Figure 3. As soil materials used in this study was autoclaved prior to testing, soil bacteria in these samples were likely introduced from the termite cuticle or from the gut during tunneling/nest building activity. Results showed a predominance of Proteobacteria and Bacteroidetes in all soil samples, although the latter was reduced in soil from the low temperature group. As observed in termite gut samples, the relative abundance of Proteobacteria was also highest in soil samples from the low temperature groups. Soil material from the high temperature group had an increased abundance of Planctomyctes and Actinobacteria, which is contrary to the decreased abundance of these phyla observed in their corresponding termite gut samples.

At the OTU level in medium temperature soil samples, the dominant OTU (00010) was classified as a deltaproteobacterium *Myxococcus* (mean relative abundance 7.6, SD= 6.9), followed by OTU00014, an unclassified member of the Chitinophagaceae family (mean relative abundance 5.5, SD=5.2). Unclassified Xanthomonadaceae, OTU00014 (mean relative abundance 5.6, SD=4.1) and OTU00076 (mean relative abundance 4.1, SD=4.4), were the most abundant in the high temperature group. In the low temperature group, OTU00001, classified to the *Enterobacter* genus (mean relative abundance 36.3, SD= 5.5), was the most dominant, followed by OTU00013, identified as a species of *Acinetobacter* (mean relative abundance 8.6, SD= 3.7). These data are in accordance with results from termite gut samples that also showed an increase in Proteobacteria *Enterobacter* (OTU00001) and *Acinetobacter* (OTU00013) in low temperature groups (Fig. 6).

## 4. Conclusions

In this study, we found shifts in environmental temperature can cause substantial changes in the microbial community of *R. flavipes* guts as well as in associated soil materials. These results suggest that termites might be more vulnerable to high heat exposure through microbial community loss/shifts, compared to low temperature. Thus, it is conceivable that termite activity might decline in warmer, southern regions (assuming similarity in gut microbial structure), and northern range edges might expand. As we are not yet able to establish a cause-effect relationship between these findings, we suggest that more studies are needed to investigate how the termite gut microbiota can modulate the host’s physiological acclimation to temperature changes and vice-versa.

## 5. Acknowledgements

We thank Avery Kuhlow for assistance in running cold tolerance assays and Rachel Slatyer who helped in the initial development of the project. We are sincerely grateful to Will Turnbull, metal worker and artist, for designing and constructing the apparatus for conducting termite cold tolerance assays. We also thank Emanuel F. Burgos-Robles for his work during the 16S rDNA library construction and the University of Wisconsin Biotechnology Center DNA Sequence Facility for providing Illumina Next Generation sequencing facilities and services. This work was funded in part through the USDA Forest Service.

## 6. Competing Interests

The authors declare no competing interests.

## Notes

### Competing Interest Statement

The authors have declared no competing interest.

